# Detecting ancient positive selection in humans using extended lineage sorting

**DOI:** 10.1101/092999

**Authors:** Stéphane Peyrégne, Michael James Boyle, Michael Dannemann, Kay Prüfer

## Abstract

Natural selection that affected modern humans early in their evolution has likely shaped some of the traits that set present-day humans apart from their closest extinct and living relatives. The ability to detect ancient natural selection in the human genome could provide insights into the molecular basis for these human-specific traits. Here, we introduce a method for detecting ancient selective sweeps by scanning for extended genomic regions where our closest extinct relatives, Neandertals and Denisovans, fall outside of the present-day human variation. Regions that are unusually long indicate the presence of lineages that reached fixation in the human population faster than expected under neutral evolution. Using simulations we show that the method is able to detect ancient events of positive selection and that it can differentiate those from background selection. Applying our method to the 1000 Genomes dataset, we find evidence for ancient selective sweeps favoring regulatory changes and present a list of genomic regions that are predicted to underlie positively selected human specific traits.

## INTRODUCTION

Modern humans differ from their closest extinct relatives, Neandertals, in several aspects, including skeletal and skull morphology (Weaver 2009), and may also differ in other traits that are not preserved in the archeological record (Laland et al. 2010; Varki et al. 2008). Natural selection may have played a role in fixing these traits on the modern human lineage. However, the selection events driving the fixation would have been restricted to a specific timeframe, extending from the split between archaic and modern humans ca. 650,000 years ago to the split of modern human populations from each other around 100,000 years ago (Prüfer et al. 2014). While methods exist, that can be used to scan the genome for the remnants of past or ongoing positive selection (Lemey et al. 2009; Nielsen et al. 2007), current methods have limited power to detect positive selection on the human lineage that acted during this older timeframe (see Sabeti et al. 2006 for a review on detection methods and their timeframes): an unusually high ratio of functional changes to non-functional changes, such as the dn/ds test, requires millions of years and often multiple events of selection to generate detectable signals (Kryazhimskiy and Plotkin 2008), while unusual patterns of genetic diversity between individuals and populations (e.g. extended homozygosity, Tajimas D, Fst) are most powerful during the selective sweep or shortly after (Oleksyk et al. 2010; Sabeti et al. 2006). A favorable substitution is not expected to leave a mark on linked neutral variation beyond 250,000 years in humans (Przeworski 2002, 2003).

The genome sequencing of archaic humans (Neandertals and Denisovans) to high coverage (Meyer et al. 2012; Prüfer et al. 2014) has spawned new methods to investigate the genetic basis of modern human traits that are not shared by the archaics (Pääbo 2014). One method, called 3P-CLR, models allele frequency changes before and after the split of two populations using the archaic genomes as an outgroup (Racimo 2016). 3P-CLR outperforms previous methods in the detection of older event of selection (up to 150,000 years ago, Figure 2 from Racimo 2016) but has little power to detect events older than 200,000 years ago in modern humans. A second method applied an approximate Bayesian computation on patterns of homozygosity and haplotype diversity around alleles that reach fixation (Racimo et al. 2014). Although, this approach expands our ability to investigate older time frames, this signal of selection also fades over time and events of positive selection older than 300kya become undetectable.

**Figure 1:**
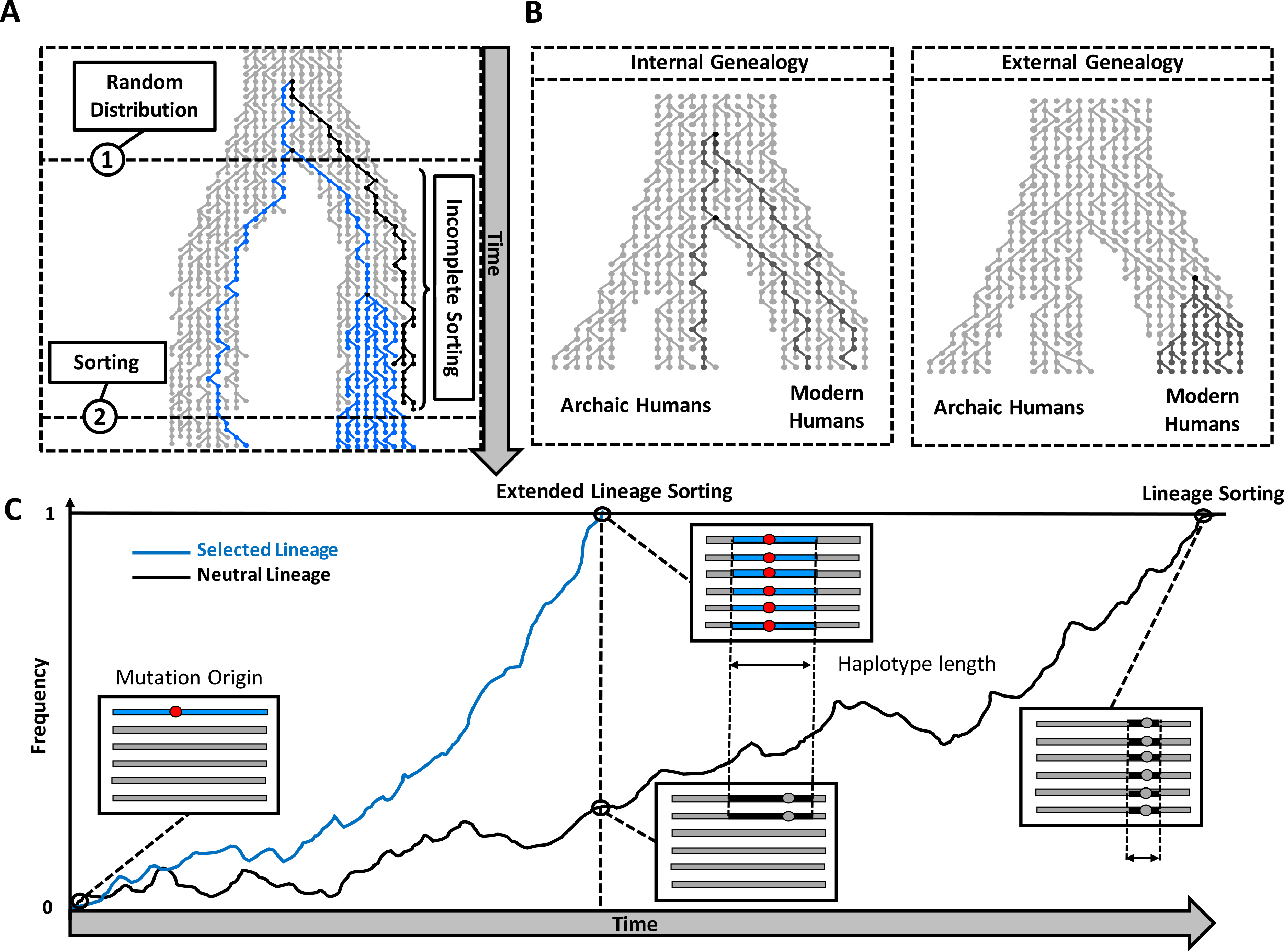
Illustration of the lineage sorting process. (A) Effects on the genealogy. The process starts with a random distribution of lineages when the ancestral population splits. The lineage in black is an out-group to lineages in blue, so that the blue lineages show a closer relationship between populations than to the black lineage (incomplete lineage sorting). When the blue lineages in the top population reach fixation (through a selective sweep for instance), any lineage from the other populations will constitute an out-group, thereby completing the sorting of lineages. (B) Two types of genealogies illustrating the possible relationships between an archaic lineage and modern human lineages. (C) Local effects in the genome at different time points. The curves represent the progression of lineage sorting for two independent regions, evolving under neutrality (black curve) and positive selection (blue curve), respectively. Longer fixation times are associated with more recombination so that neutrality produces smaller external regions.

**Figure 2:**
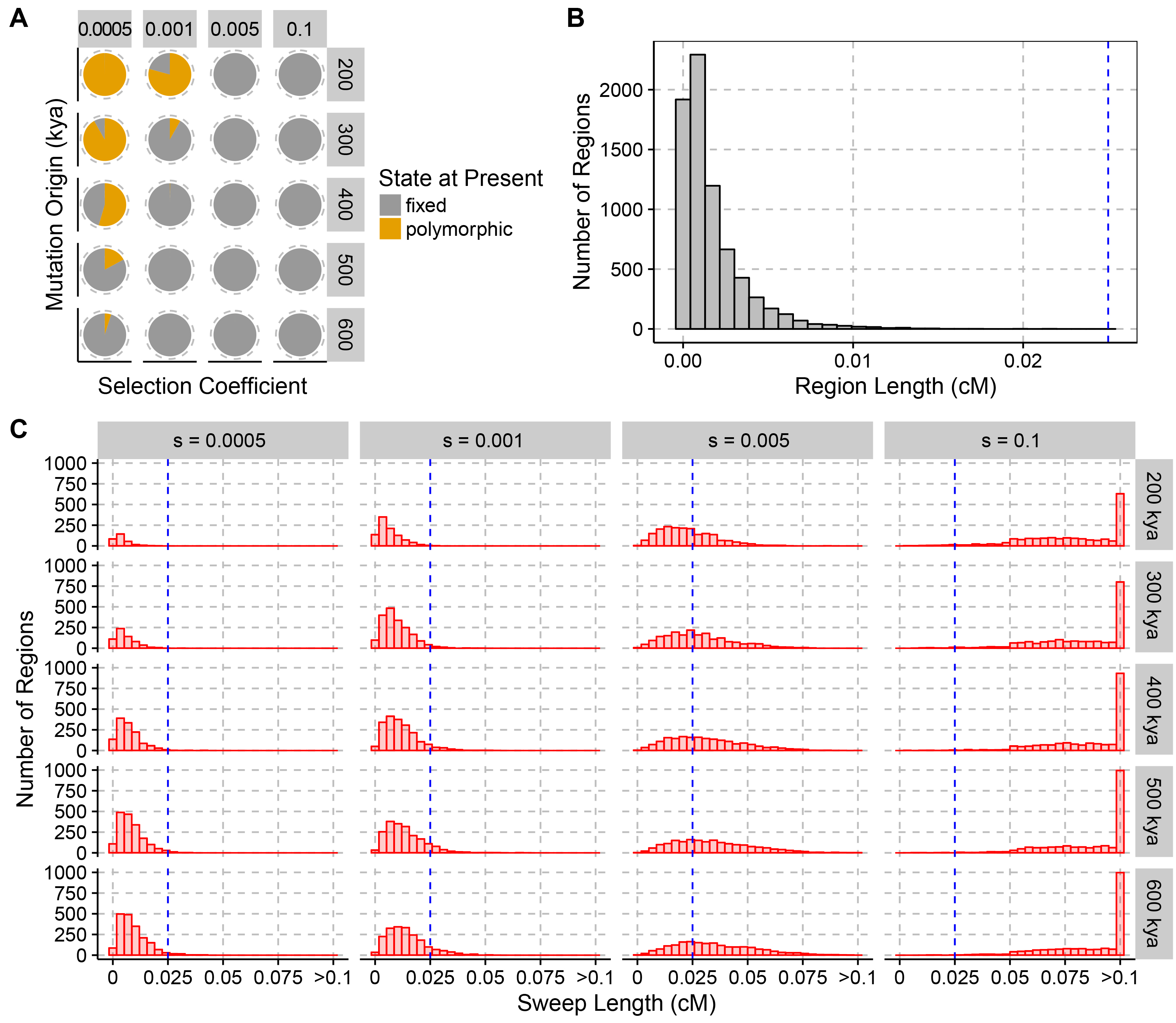
(A) Fraction of selected alleles reaching fixation (grey) or segregating (orange) at present, depending on the strength of selection (columns) and the age of the mutation (rows, in kya) in our simulations. Events for which the selected variant was lost are not shown. (B) Distribution of the genetic length of external regions simulated under neutrality. (C) Distributions of the genetic length of external regions depending on the strength of selection (columns) and age of mutations in kya (rows). The blue line corresponds to the upper limit for the length of external regions produced under neutrality from (B).

Based on a method introduced by Green et al. (2010), Prüfer et al. (2014) presented a hidden Markov model that identifies regions in the genome where the Neandertal and Denisovan individuals fall outside of present-day human variation (i.e. the archaic lineages fall basal compared to all present-day humans), and applied the model to detect selective sweeps on the modern human lineage. Regions that are unusually long are candidates for ancient selective sweeps as variants are likely to have swept rapidly to fixation, dragging along with them large parts of the chromosomes that did not have time to be broken up by recombination. While this method is, in principle, expected to be able to detect events as old as the modern human split from Neandertals and Denisovans, this power was never formally tested and it has several other shortcomings. First, the method was limited to modern human polymorphisms, ignoring the additional information given by fixed substitutions. Second, the method does not fit parameters to the data, but requires these parameters to be estimated through coalescent simulations.

Here, we introduce a refined version of this method, called ELS (Extended Lineage Sorting), that models explicitly the longer regions produced under selection, and includes the fixed differences between archaic and modern human genomes as an additional source of information. The ELS method also takes advantage of an Expectation-Maximization algorithm to estimate the model parameters from the data itself, making it free from assumptions regarding human demographic history.

To evaluate the power of the ELS method to detect ancient selective sweeps we tested its performance under scenarios of background selection and neutrality. Finally, we present an updated list of candidate regions that likely underwent positive selection on the modern human lineage since the split from the common ancestor with Neandertals and Denisovans.

## RESULTS

### Selection causes extended lineage sorting between closely related populations

The ancestors of modern humans split from the ancestors of Neandertals and Denisovans between 450,000 and 750,000 years ago (Prüfer et al. 2014). Because the two newly formed descendant groups sampled the genetic variation from the ancestral population, a derived variant can be shared between some members of both groups, while other individuals show the ancestral variant. At these positions, some lineages from one group share a more recent common ancestor with some lineages in the other group than within the same group (Rosenberg 2002), a phenomenon called incomplete lineage sorting (Figure 1A).

Eventually, a derived allele may reach fixation as part of a region that has not been unlinked by recombination. In these regions all descendants will derive from one common ancestor and any lineage from the other population will constitute an out-group, i.e. all lineages are sorted. Because of recombination, the human genome is a mosaic of independent evolutionary histories and the process of lineage sorting is expected to randomly affect regions, until, ultimately, all lineages will be sorted. In the case of modern humans, only a fraction of the regions in the genome are expected to show lineage sorting (Prüfer et al. 2014), and the genome can be partitioned into regions where an archaic lineage falls either within the variation of modern humans (internal region) or outside of the human variation (external region) (Figure 1B).

While lineage sorting can occur under neutrality, selection on the modern human branch is expected to always lead to external regions as long as the selective sweep finished. In cases where the selective sweep is sufficiently strong, there will not be sufficient time for recombination to break the linkage with neighboring sites and a large region will reach fixation (extended lineage sorting, ELS, Figure 1C). In contrast, selection on standing variation may fail to generate such large regions, since recombination can act on the haplotype(s) with the prospective advantageous variant before selection sets in. We note that neither demography nor selection on the archaic lineage affect the lineage sorting within modern humans and thus the power to detect selective sweeps.

### Expected Incomplete Lineage Sorting among Humans to Archaics

We used coalescent simulations to determine the incidence and expected length of regions resulting from incomplete lineage sorting in modern humans. Using a model of human demographic history (Yang et al. 2014), we estimated the fraction of lineage sorting in modern humans in regards to Neandertals and Denisovans. In simulations with 370 African chromosomes, and assuming a uniform recombination rate, about 10% of the archaic genome is more divergent than the time to the most recent common ancestor of all sampled human variation. The length of the external regions is expected to be about 0.0016 cM (95%-CI: 0.001-0.0095 cM; e.g. 1-9.5kb for a recombination rate of 1cM/Mb) with the longest regions in the order of 0.02 cM. In contrast, internal regions are expected to be 0.012 cM long (95%-CI: 0.0097-0.07 cM).

### Minimum Strength of Selection to Produce Detectable Sweep Signals

We investigated the range of selection coefficients that could have led to the fixation of a lineage after the split with the Archaic hominins, but before the differentiation of genetically modern humans about 100−120 kyr ago (Li and Durbin 2011) by simulating mutations occurring at different times and evolving with different selection coefficients. While the simulations show that completed selective sweeps could have occurred with selection coefficients as low as 0.0005 (Figure 2A), the length distribution of haplotypes reaching fixation is indistinguishable from neutrality for selection coefficients under 0.001 (Figure 2, B and C). Under neutrality, the average length of external regions was 0.02 cM and remained below 0.03cM for most simulations with a selection coefficient of 0.001. In contrast, external regions longer than 0.1cM were observed for selection coefficients above 0.05. Therefore, detectable signals are expected to be biased towards strong events with a selection coefficient larger than 0.001.

### Hidden Markov Model to Detect Extended Lineage Sorting

To detect regions of Extended Lineage Sorting, we modeled the changes of local genealogies along the genome with a hidden Markov model. We distinguish two types of genealogies, internal or external, depending on whether the archaic lineage falls inside or outside of the human variation respectively (Figure 3A). The model includes a third state corresponding to extended lineage sorting, and external regions produced by this state are required to be longer, on average, than those produced by the external state. The three states are inferred from the state of the archaic allele (ancestral or derived) either at a polymorphic position in modern humans or at a position where modern humans carry a fixed derived variant. In the following, we describe the different statistical properties expected for each type of genealogy.

**Figure 3:**
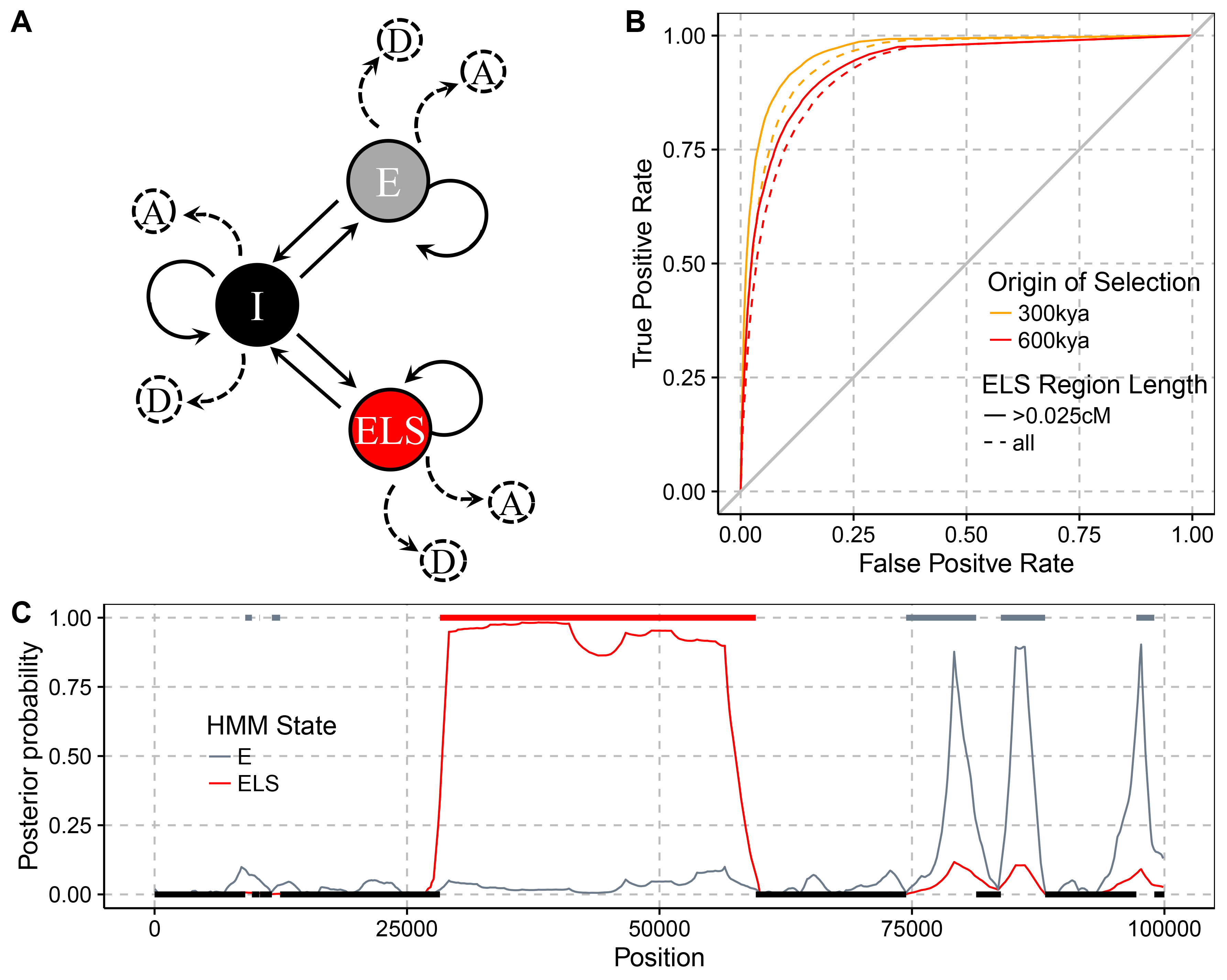
(A) Graphical representation of the Extended Lineage Sorting Hidden Markov Model. States are depicted by nodes and transitions by edges. Each state emits an archaic allele as either derived, D, or ancestral, A, depending on the type of site in the modern human population (fixed or segregating at a given frequency). States are labelled I for Internal, E for External and ELS for Extended Lineage Sorting. (B) Receiver Operator Curves for varying cutoffs on the posterior probability of the ELS state and counting the number of sites in ELS regions that were correctly labeled. All bases labelled ELS outside of simulated ELS regions are considered false positives. Sites in ELS regions with a posterior probability below the cutoff are considered false negatives. (C) Example of the labelling of a simulated ELS region. Horizontal bars indicate true external (top) and internal (bottom) regions. The posterior probability is shown in red for ELS regions and in grey for E regions. The region overlapping position 50,000 (red bar) is caused by a simulated selective sweep.

We first consider external regions. At modern human polymorphic sites, the archaic genome is expected to carry the ancestral variant since the derived variant would indicate incomplete lineage sorting. To account for sequencing errors or misassignment of the ancestral state, we allow a probability of 0.01 for carrying the derived allele (see Material and Methods). At sites where the derived allele is fixed, the archaic genome will often carry the derived state, if the fixation event occurred before the split of the archaic from the modern human lineage, or, occasionally, the ancestral state, if the fixation event is more recent and occurred after the split.

For internal regions, the archaic is expected to share the derived allele at modern human fixed derived sites, but can carry the ancestral allele in our model to accommodate errors, albeit with low probability. In contrast, at sites that are polymorphic in modern humans, the probabilities of observing the ancestral or the derived allele in the archaic genome will depend on the age of the derived variant, with young variants being less likely to be shared compared to older variants. The frequency of the derived variant in the modern human population can be used as a proxy for its age and the emission probabilities in our model take the modern human derived allele frequency into account (see Material and Methods).

We modeled the transition probabilities between internal and external regions (related to the length of the regions) by exponential distributions. The extended lineage sorting state has the same chance of emitting derived alleles as the other external state but is required to have a larger average length. We used the Baum-Welch algorithm (Durbin et al. 1998), an Expectation-Maximization algorithm, to estimate the emission probabilities, and estimate the transition probabilities with a likelihood maximization algorithm.

### Accuracy of Parameter Estimates and Inferred Genealogies

We first investigated the performance of the parameter inference on simulated data under neutral evolution. We found that the estimated probabilities for encountering ancestral/derived alleles in external and internal regions fit the simulated parameters well (on average less than ± 0.08 from simulated under all tested conditions) (Supplemental Figures S1 and S2), while the estimated length of internal and external regions deviate more from the simulated lengths (around 15% overestimate of the mean length, Supplemental Figure S3). However, we found that the model exhibits better accuracy in labelling the correct genealogies with the estimated length parameters compared to the simulated true values (Supplemental Figure S4). This difference seems to originate from the difficulty in accurately detecting very short external regions or internal regions with very few informative sites. We note that detecting selection is not affected by this problem since we are primarily interested in detecting long external regions. Including fixed differences improves the power to assign the correct genealogies compared to a version of the method without this additional source of information (Supplemental Figure S4).

We do not expect ELS regions to be detected in our neutral simulations and indeed we found that either the estimated proportion of ELS converged to zero or the maximum likelihood estimate for the length of ELS and external regions converge to the same value (49% and 51% of simulations respectively). A likelihood ratio test comparing a model without the ELS state to the full model with the ELS state also showed no significant improvement with the additional state in almost all neutral simulations (only one likelihood ratio test out of 100 simulations showed a significant improvement after Bonferroni correction for multiple testing).

We then evaluated the accuracy of the ELS method to assign the correct genealogy to regions based on sequences obtained through coalescent simulations with selection (Figure 3, B and C). In these simulations, the underlying genealogy at each site along the sequences is known and can be compared to the estimates. To be conservative, we only focus on results with the smallest selection coefficient (s=0.005) that produces regions long enough to be detectable. In Figure 3B we show the accuracy for labelling the extended lineage sorting regions dependent on the posterior probability cutoff for the ELS state. The results demonstrate that the model has sufficient power to accurately label sites that experienced selection with a coefficient s>=0.005 and an occurrence of the beneficial mutation as long as 600,000 years ago.

We also used the simulations of positive selection events (s=0.005) with two different times at which the beneficial mutation occurred, 300kya and 600kya, to test how often the beneficial simulated variant fall within a detected ELS region (Supplemental Table S1). To put this rate of true positives into perspective, we also counted how many ELS regions did not overlap the selected variant (false positives). A large fraction of selected mutations were detected (87-92%). However, we also found a substantial fraction of false positive ELS regions (10-11%). When restricting detected ELS regions to those that are longer than 0.025cM, we find less than 0.1% false positives compared to 65-68% true positives. Not all simulated regions with a selection coefficient of 0.005 produce ELS regions of this size, so that the rate of true positives for truly long regions is expected to be higher. For all following analysis, we used this minimal length cutoff of 0.025 cM.

### Role of Background selection

Background selection is defined as the constant removal of neutral alleles due to linked deleterious mutations (Charlesworth et al. 1993). In regions of the genome that undergo background selection, a fraction of the population will not contribute to subsequent generations, causing a reduced effective population size. As a consequence, remaining neutral alleles can reach fixation faster than under neutrality, potentially producing unusually long external regions that could be mistaken as signals of positive selection. We investigated the effects of background selection by running forward simulations with parameters that mimic the strength and extent of background selection estimated for the human genome (Messer 2013). While background selection simulations did produce some long outlier regions that fall outside the distribution observed in neutral simulations, most regions are still smaller than regions simulated with positive selection at a conservative selection coefficient of 0.005 (Figure 4A). Indeed, among the 1160 external regions detected in our simulations of background selection (s=0.05, Figure 4A) only six were labeled as ELS and only three passed the minimal length filter of 0.025 cM.

**Figure 4:**
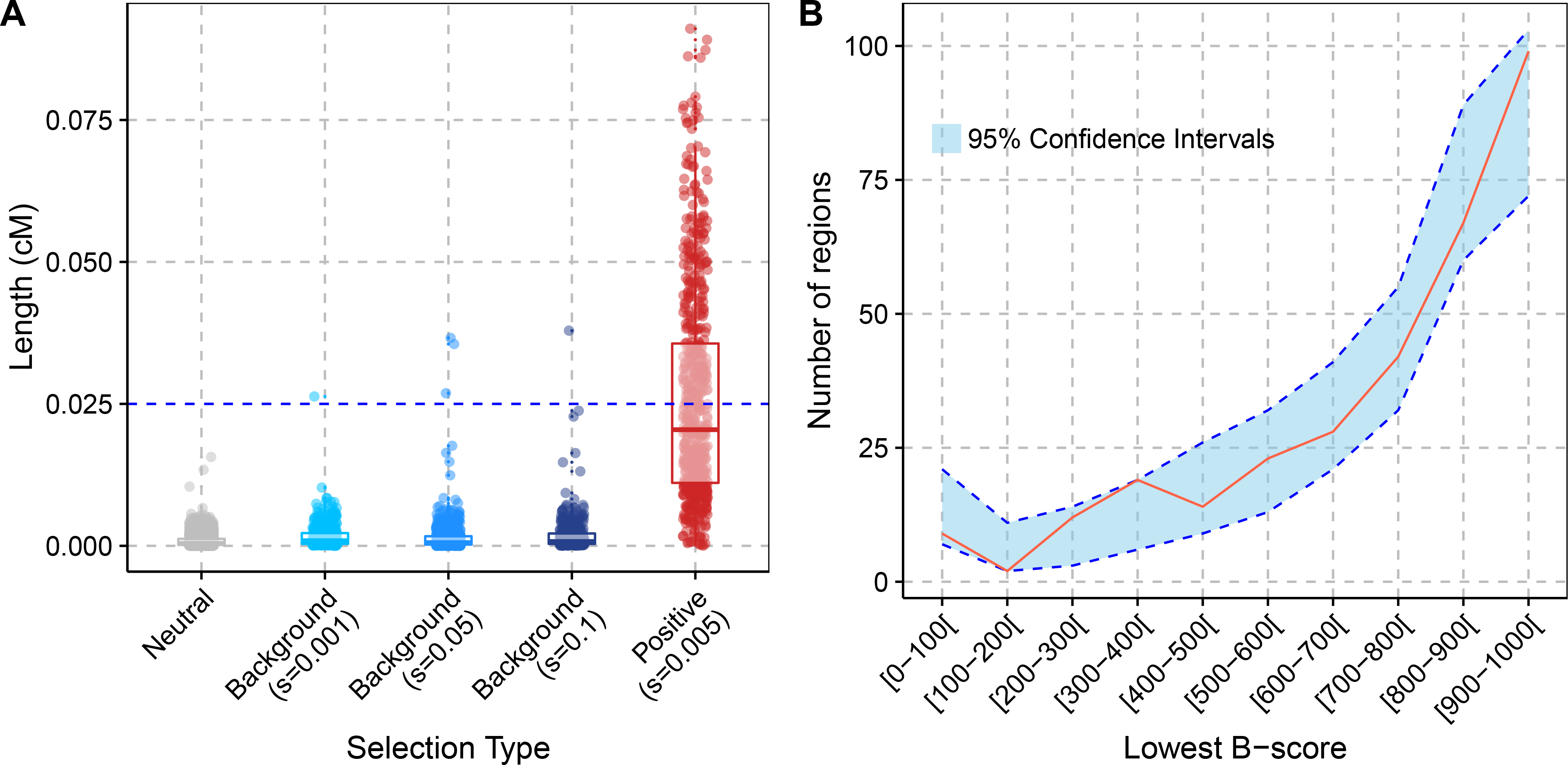
Effects of background selection. (A) Comparison of the length of ELS regions in simulations of different scenarios. For the distribution under background selection, the s parameter corresponds to the average selection coefficient from the gamma distributio n (shape parameter of 0.2). We assumed that the deleterious mutations are recessive with dominance coefficient h=0.1. The horizontal blue line corresponds to the length cutoff applied to the real data. (B) Distribution of B-scores in the candidate sweep regions (red curve) compared to sets of random regions with matching physical lengths (blue area with dotted blue lines indicating the 95% confidence intervals over 1000 random sets of regions). The lowest B-score (i.e. stronger background selection) was chosen when a region overlapped several B-score annotations.

### Candidate Regions of Positive Selection on the Human Lineage

To identify ancient events of positive selection on the human lineage, we applied the ELS method to African genomes from the 1000 genomes project (Abecasis et al. 2012). We disregarded non-African populations since Neandertal introgression in these populations could mask selective sweeps and lead to false negatives. A model with ELS fits the data significantly better than a model without the ELS state for all chromosomes and for both tested recombination maps (p-value < 1e-8, Supplemental Table S2).

We identified 81 regions of human extended lineage sorting for which both recombination maps support a genetic length greater than 0.025cM (average length: 0.05 cM). Depending on the recombination map, the longest overlap between the maps is 0.12 (African-American map) or 0.17 (deCode map) cM long, which is three to four times longer than the longest regions produced under background selection in our simulations. An additional 233 regions are longer than 0.025cM according to only one recombination map, with 71% of those additional regions showing support for the ELS state using both recombination maps. This suggests that the variation in the candidate set mostly stems from uncertainty about recombination rates. We will refer to the set of 81 regions as the core set (Supplemental File S1) and the set including the 233 putatively selected regions found with just one recombination map as the extended set (314 regions, Supplemental File S2).

For completeness, we also ran our model on the X chromosome and identified 12 additional candidates (43 if we consider candidates found with at least one recombination map), applying a more stringent length cutoff of 0.035 cM to account for the stronger effects of random drift on this chromosome (cf. Material and Methods). Interestingly, we also found a significant increase of posterior probabilities for selection within previously reported regions under potential recurrent selective sweeps in apes (Dutheil et al. 2015; Nam et al. 2015) (*Mann-Whitney U* one-sided test, *P*-value < 2.2e-16, Supplemental Table S3).

The detected selection candidate regions on the autosomes do not show a decrease in B scores (McVicker et al. 2009), a local measure of background selection strength, compared with random regions (Figure 4B; Wilcoxon rank sum test comparing the average B-scores with permuted regions, *P*-value=0.565, or comparing the lowest B-scores in our regions to permuted regions, *P*-value=0.504). This suggests that candidate regions are not primarily generated by strong background selection.

We compared our candidate regions to the top candidates of 8 previous scans for selection, including iHS, Fst, XP-CLR and HKA (Cagan et al. 2016; Chen et al. 2010; Hudson et al. 1987; Malécot 1948; Pybus et al. 2014; Voight et al. 2006; Wright 1951). Using the estimated TMRCA among Africans for each identified region/site, we found that our ELS scan identified significantly older events than other screens (Figure 5, *Mann-Whitney U* tests, Supplemental Table S4). We found 23 regions from the core set (detected by both recombination maps) overlapping with candidates from previous scans and 68 for the extended set (detected by at least one recombination map); neither overlap is more than expected at random (*P*-values are 0.06 and 0.595 respectively). In contrast, our candidate regions overlap more often candidate regions from 3P-CLR (Racimo 2016) and the ABC approach for detecting ancient selection (Racimo et al. 2014) than expected by chance (*P*-values<0.05; Supplemental Table S5).

**Figure 5:**
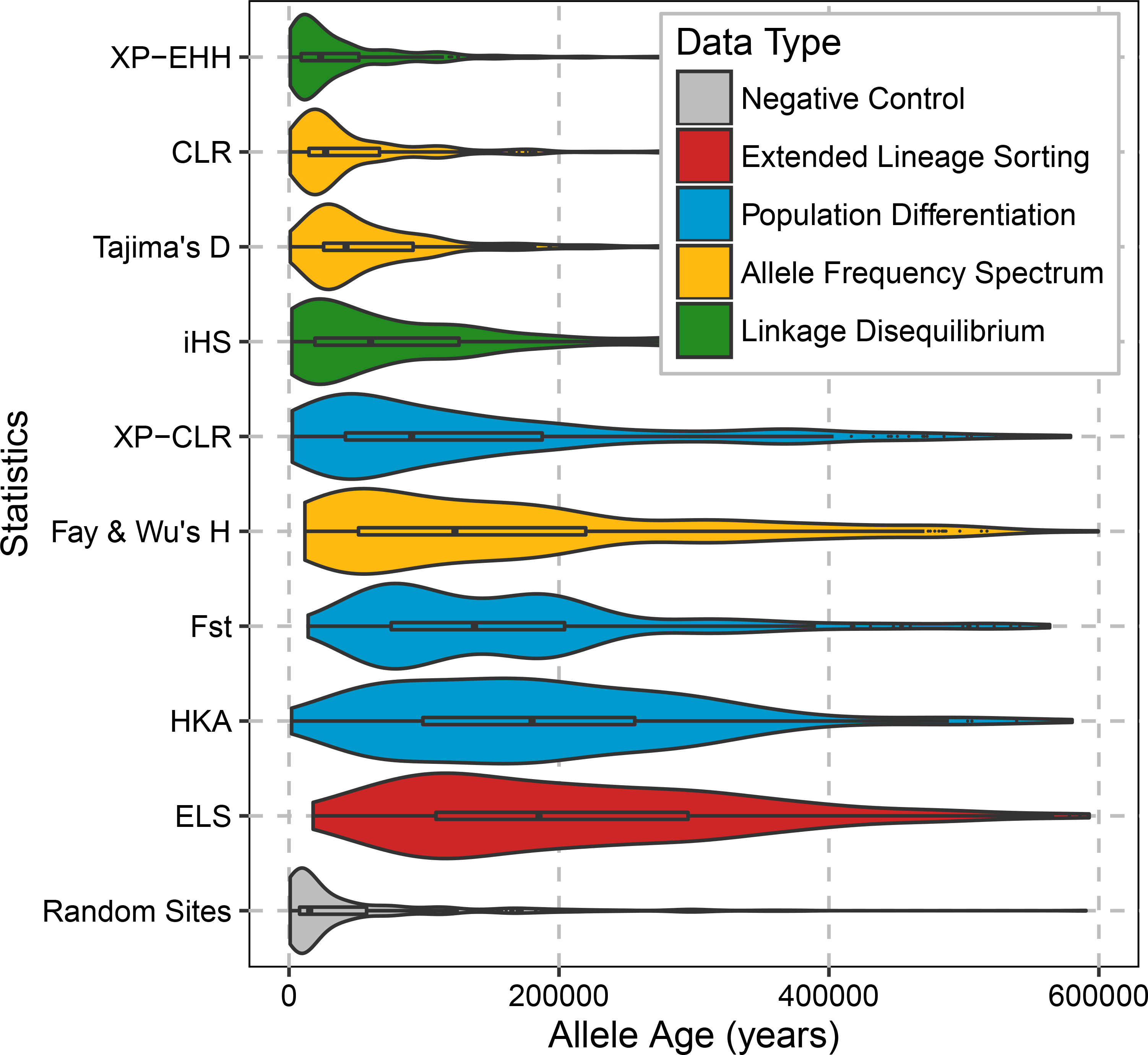
Distributions of estimated ages of the modern human segregating derived variants with the highest frequency in putatively selected regions or the age of the derived variants at sites identified by various genome-wide scans. Our candidate regions are labelled as ELS, for Extended Lineage Sorting, other candidate regions are from (Cagan et al. 2016; Pybus et al. 2014). The color coding indicates the type of signal detected by each method. Ages were estimated by ARGweaver (Rasmussen et al. 2014). We only report events between 0 and 600kya.

### Biological functions of the candidate regions

Since positive selection acts on advantageous phenotypes that are caused by changes to functional elements in the genome, we would expect that our candidate regions overlap functional elements in the genome more often than expected.

We first tested this hypothesis by counting the overlap between sweep candidate regions and protein coding genes (Ensembl release 82). We find no statistically significant overlap of ELS regions with protein coding genes compared to randomly placed regions of the same size (*P*-value = 0.671 and 0.124, for core and extended set, respectively; Figure 6A). Previous work has identified 96 proteins that carry human fixed derived non-synonymous changes compared to Neandertal and Denisova, which constitute a particularly interesting subset of potentially functional changes to genes that may have been caused by selective sweeps (Prüfer et al. 2014). We found no overlap between these genes and the core set of sweep candidate regions that were identified by both recombination maps. However, when considering the extended set of sweep candidate regions, 11 regions overlapped such genes: *ADSL, BBIP1, ENTHD1, HERC5, KATNA1, KIF18A, NCOA6, PRDM10, SCAP, SLITRK1* and *ZNHIT2*. This overlap is significantly larger than expected by chance (only 2 genes are expected on average; *P*-value < 10^−3^). In all instances, the candidate regions contained at least one fixed amino acid change. Since fixed changes are part of the information used to infer external regions, it stands to reason that the presence of such a change may bias towards observing an overlap with candidate regions (72/81 core regions and 275/314 regions from the extended set contain fixed changes). However, we note that the overlap with fixed amino acid changes is also significantly larger than the overlap with other fixed changes (963 of 20347 fixed changes fall within candidate regions from the extended set; binomial *P*-value=0.006).

**Figure 6:**
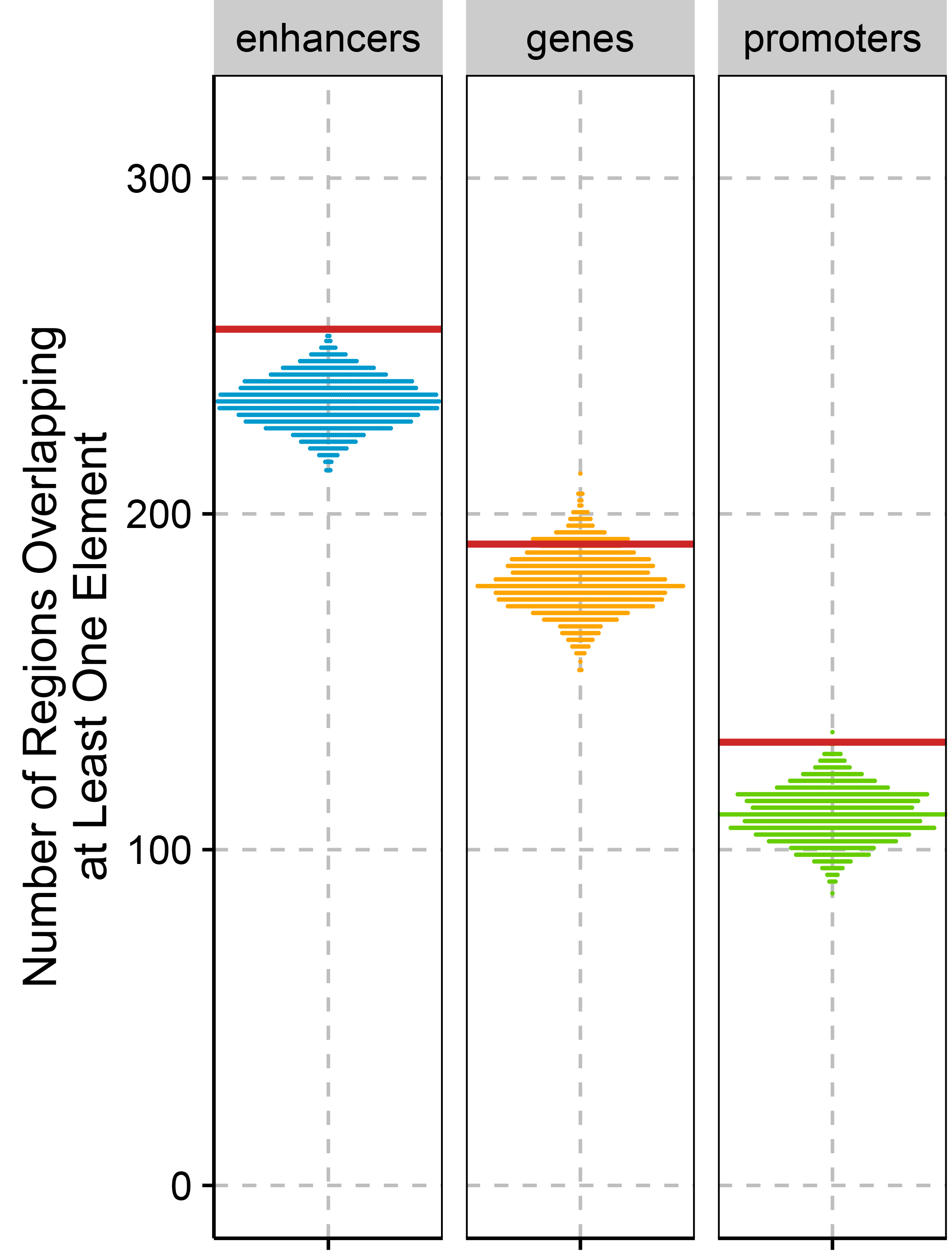
Enrichment for regulatory elements (enhancers, *P*-value<0.001, protein-coding genes, *P*-value=0.124, and promoters, *P*-value=0.002) in the extended set of 314 candidate sweep regions. The distributions were obtained by randomly placing candidate regions in the genome to obtain lists of regions with similar physical length. The red lines represent the value observed in the real extended set.

Phenotype may also be influenced by regulatory changes that affect gene expressions. Interestingly, we found a significant enrichment for regions overlapping enhancers and promoters (*P*-value<0.001 and *P*-value=0.002, respectively; see Figure 6A) when considering the extended set of 314 candidate regions. However, this enrichment was not significant for the smaller core set of candidates.

To further investigate the biological function of our regions, we tested for gene ontology enrichment in genes within the extended set of regions. No category showed significant enrichment when comparing to randomly placed regions of identical sizes in the genome (Material and Methods). We also assigned genes that overlap our extended dataset to tissues in which they show the significantly highest expression and found again no enrichment. In an attempt to include potential regulatory changes in the enrichment test, we assigned genes to candidate regions when a region fell upstream or downstream of a gene (see Material and Methods). Although many candidate genes that were annotated in this way were expressed highest in the brain or the heart (OR=2.10 for both tissues), this enrichment is not significant when correcting for gene length and multiple testing (FWER=0.336 and 0.997 respectively, Supplemental Table S7).

Additional work will be required to investigate the phenotypic consequences of changes in candidate regions for selection. To facilitate this work, we provide an annotated list of fixed or nearly fixed sites on the human lineage that fall within our candidate regions (Supplemental File S3).

### Overlap with Neandertal Introgression

Introgression from Neandertals and Denisovans into modern humans occurred approximately 37,000 to 86,000 years ago (Fu et al. 2014, 2015; Sankararaman et al. 2012, 2016). For those advantageous derived variants that arose on the modern human lineage prior to introgression, we would expect that selection may have acted against the re-introduction of the ancestral variant through admixture. We tested whether this selection may have affected the distribution of Neandertal introgressed DNA around fixed changes in candidate sweep regions. Out of a total of 963 fixed derived variants in Africans overlapping the extended set of sweep regions, 240 (25%) show the ancestral allele in non-Africans and show evidence for re-introduction by admixture using a map of Neandertal introgression (Vernot and Akey 2014). This level of Neandertal ancestry is comparable to the genome-wide fraction of out-of-Africa ancestral alleles at African fixed derived sites (∼26%; bootstrap *P*-value=0.583). We also find no significant reduction in frequency of Neandertal ancestry around candidate substitutions in sweep regions, when comparing one randomly sampled fixed African substitution per region against random regions matched for size and distance to genes (Supplemental Figure S6 and S7).

If selection against the re-introduction of an ancestral variant were very strong, selection may have depleted Neandertal ancestry in a large region surrounding the selected allele. Interestingly we find some of our sweep candidate regions that fall within the longest deserts of both Neandertal and Denisova ancestry (Table 1) (Vernot et al. 2016). A significantly high number of the core set of regions fall in these deserts (5/81 regions, *P*-value=0.024), while the extended set shows no significant enrichment (9/314 regions, *P*-value=0.205).

**Table 1:** Genes from the core set of candidate regions overlapping with long deserts of Neandertal and Denisovan ancestry.

| Chromosome | Start | End | Overlapping Genes | Overlapping Regulatory Domains |
| --- | --- | --- | --- | --- |
| chr1 | 104000000 | 104154236 | *AMY2B, RNPC3* | *COL11A1* |
| chr1 | 113429666 | 113560554 | *SLC16A1* | *FAM19A3, LRIG2* |
| chr3 | 77027850 | 77033270 | *ROBO2* | - |
| chr7 | 122320038 | 122379695 | *RNF133, RNF148, CADPS2* | *TAS2R16* |
| chr10 | 107809941 | 107866217 | - | SORCS1, SORCS3 |

## DISCUSSION

Many genetic changes set modern humans apart from Neandertals and Denisovans but their functions remain elusive. Most of these changes probably resulted in either no change to the phenotype or to a selectively neutral change. However, in rare instances selection may have favored changes modifying the appearance, behavior and abilities of present-day humans. Unfortunately, current methods to identify selection have limited power to detect such old events of positive selection (Przeworski 2002, 2003; Sabeti et al. 2006).

Here, we introduce a hidden Markov model to detect ancient selective sweeps based on a signal of extended lineage sorting. Using simulations we were able to show that the method can detect older events of selection as long as the selected variant was sufficiently advantageous. The power to detect older events is due to the fact that the method increases in power with the number of mutations that accumulated after the sweep finished. We also showed that background selection can cause false signals and have chosen a minimum length cutoff on candidate regions. While this cutoff reduces the number of false positives due to background selection, we note that this cutoff is expected to exclude *bona fide* events of positive selection, too.

We applied the ELS method to 185 African genomes, the Altai Neandertal genome and the Denisovan genome, and detected 81 candidate regions of selection when requiring a minimum genetic length supported by two independent recombination maps. The uncertainty in the recombination maps has a large effect on our results, as shown by the much larger number of 314 regions identified by either recombination map. Recombination rates over the genome are known to evolve rapidly (Lesecque et al. 2014) and of particular concern are recent changes in recombination rates that make some regions appear larger in genetic length than they were in the past. By comparing the current recombination rates in our regions to recombination rates in the ancestral population of both chimpanzee and humans (Munch et al. 2014), we identified some candidate regions that may have increased in recombination rates (Supplemental Table S7). However, it is currently impossible to date the change in recombination rates confidently and these candidate sweeps may post-date the change.

A particular strength of our screen for selective sweeps is the ability to detect older events, as indicated by the estimated power to detect simulated events of positive selection of old age and moderate strength. This sets the ELS method apart from previous approaches that made use of archaic genomes, which were geared towards detecting younger events with an age of less than 300,000 years ago (Racimo 2016; Racimo et al. 2014). Despite this difference, we found significant overlap between the ELS candidates and the candidates identified by these other approaches, while the overlap with other types of positive selection scans is smaller. Among our candidates, 55 are novel candidates (234 if considering the extended set) that were not detected in any of the previous screens, including previous versions of the screen without fixed differences (Supplemental Figure S5).

While we find no difference in the fraction of genes in selected regions compared to randomly placed regions, we detect an enrichment for enhancers and promoter regions. This result is in agreement with the hypothesis that regulatory changes may play an important role in human-specific phenotypes (Carroll 2003; Enard et al. 2014; King and Wilson 1975), maybe more so than amino-acid changes (Hernandez et al. 2011; see also Enard et al. 2014 and Racimo et al. 2014). Interestingly, several gene candidates falling within sweep regions play a role in the function and development of the brain. A particularly interesting observation is the potential selection on both the ligand *SLIT2* and its receptor *ROBO2*, which reside on chromosome 4 and 3 respectively (see Supplemental File S3 for an annotated list of changes in those genes). Members of the Roundabout (ROBO) gene family play an important role in guiding developing axons in the nervous system through interactions with the ligands SLITs. SLITs proteins act as attractive or repulsive signals for axons expressing different ROBO receptors. *ROBO2* has been further associated with vocabulary growth (St Pourcain et al. 2014), autism (Suda et al. 2011), and dyslexia (Fisher and DeFries 2002) and is involved in the development of neural circuits related to vocal learning in birds (Wang et al. 2015). Interestingly, *ROBO2* is also in a long desert of both Denisovan and Neandertal ancestry in non-Africans.

We also identified interesting brain-related candidates on the X chromosome, among them *DCX*, a protein controlling neuronal migration by regulating the organization and stability of microtubules (Gleeson et al. 1999). Mutations in this gene can have consequences for the expansion and folding of the cerebral cortex, leading to the “double cortex” syndrome in females and “smooth brain” syndrome in males (Gleeson et al. 1998).

We have presented a new approach to detect ancient selective sweeps based on a signal of extended lineage sorting. Applying this approach to modern human data revealed that selection may have acted primarily on regulatory changes. With population level sequencing of non-human species becoming more readily available we anticipate that this approach will help to reveal the targets of ancient selection in other species

## MATERIALS AND METHODS

### Data

We used 185 unrelated Luhya and Yoruba individuals from the 1000 Genomes Project phase I (Abecasis et al. 2012), corresponding to 370 sets of autosomes and 279 X chromosomes. From this dataset, we extracted allele counts at single nucleotide polymorphism (SNP) sites using vcftools (Danecek et al. 2011). In order to add sites where all Africans differ from the common ancestor with chimpanzee, we first compiled a list of all sites where six high-coverage African genomes (Mbuti, San and Yoruban A and B-panel individuals from Prüfer et al. 2014) are identical. A site was regarded fixed different when the whole genome alignments of at least three out of four ape reference genome assemblies (chimpanzee (panTro3), bonobo (panPan1.1), gorilla (gorGor3) and orangutan (ponAbe2); lastz alignments to the human genome GRCh37/hg19 prepared in-house and by the UCSC genome browser (Speir et al. 2016)) had coverage and were different from the African allele, and when the site was not marked as polymorphic among the 1000 Genomes Luhya and Yoruba individuals.

Neandertal and Denisova alleles at polymorphic and fixed positions were extracted from published VCFs and positions were further filtered to sites passing the published map35_100 filter for both the Denisova and Neandertal genotypes (Prüfer et al. 2014). Sites where either Neandertal or Denisova carried a third allele were disregarded.

Over all autosomes, 11 million SNPs passed the filters in addition to 6.6 million African fixed variants. For the X chromosome, pseudoautosomal regions, defined as chrX: 60,001-2,699,520, chrX: 154,931,044-155,260,560 in hg19 coordinates (http://www.ncbi.nlm.nih.gov/assembly/2758/), were filtered out and around 315,000 SNPs as well as 248,000 African fixed variants remained for analysis.

Genetic distances between those positions were calculated using the African-American (Hinch et al. 2011) and the DeCode (Kong et al. 2010) recombination maps (available in Build 37 from http://www.well.ox.ac.uk/∼anjali/). Both maps were chosen since they estimate recombination rates from events that occurred within a few generations before present. Recombination maps based on older events (i.e. LD based map) can underestimate recombination rates in regions that underwent recent selective sweeps, potentially masking true signals.

Changes of recombination rates along the human lineage could also limit our power to detect selected regions, and we used an ancestral recombination map of the human-chimpanzee ancestor to annotate top candidate regions (Supplemental Table S7) (Munch et al. 2014).

### Hidden Markov model

We would like to estimate for each informative position the probabilities for the three possible genealogies external (*E*), internal (*I*) and extended lineage sorting (*ELS*) given the observed data. Formally, and following the notation from Durbin et al. 1998, we calculate *P*(*π*_*i*_ = *k*|*x*) where *i* denotes the position, *k* ∈ {*E, I, ELS*} and *x* the sequence of observations with the *i*th observation denoted *x_i_*. With the genetic distance *d* between consecutive sites and *l*_*k*_, the average genetic length of a region in state *k*, we specify the transition probabilities between identical states as 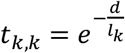. Transitions from *I* to the states *ELS* and *E* depend on an additional parameter *p*, the proportion of transitions from *I* to *ELS*, and their probability is given by 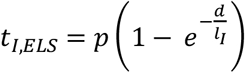 and 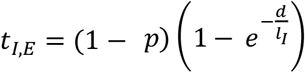. Lastly, transitions from the two external states to internal have the probability 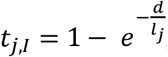, with *j* ∈ {*E, ELS*}. By construction, transitions between E and ELS genealogies are not allowed: it would not be possible to detect such transitions as those two states have the same statistical properties.

The inference further requires the probability for observing an ancestral or derived allele in the archaic at a site *i* with a derived allele frequency *f*_*i*_ > 0 in modern humans (noted *x*_*i*_) given that the true genealogy is *k* ∈ {*I, E, ELS*}: *e*_*k*_(*x*_*i*_) = *P*(*x*_*i*_||*π*_*i*_ = *k*). We assume that ∀*x*: *e*_*ELS*_(*x*) = *e*_*E*_(*x*), i.e. that both external states give rise to ancestral and derived alleles in the archaic with equal probabilities given the same observation. Since external regions are not expected to give rise to derived sites when the derived allele is segregating in modern humans, the only sources for such an observation can be errors or independent coinciding identical mutations and we define an error rate for external regions: *ϵ*_*E*_ = *e*_*E*_ (*x*_*i*_ = *derived*, *f*_*i*_ < 1). Similarly fixed derived sites are expected to show the derived allele in the archaics if the local genealogy is internal and we define an error rate for internal regions: *ϵ*_*I*_ = *e*_*I*_ (*x*_*i*_ = *derived*, *f*_*i*_ = 1).

We compute the posterior probability *P*(*π*_*i*_ = *k* | *x*) that an observation *x*_*i*_ came from state *k* given the observed sequence *x* as: 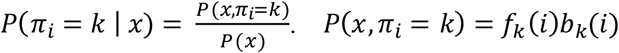 where 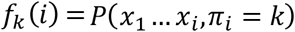 and 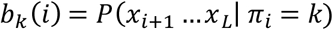 are the output of the Forward and Backward algorithms respectively (Durbin et al. 1998; Rabiner 1989). *P*(*x*) corresponds to the likelihood of the data given our model and was also calculated from the Forward algorithm.

### Parameter estimate

We used the Baum-Welch algorithm to estimate all emission probabilities with the exception of *ϵ*_*E*_, the proportion of segregating sites derived in the archaic genome in external regions, due to limited accuracy in the estimates. We set this last parameter to a value of 0.01, a conservative upper limit on contamination and sequencing error in the two high-coverage archaic genomes. The Baum-Welch algorithm was run for a maximum of 40 iterations and the convergence criteria was set to a log-likelhood maxima difference of less than 10^−4^.

We estimated the remaining parameters (average lengths of regions and the proportion of transitions to the ELS state) using the derivative free optimization method COBYLA (Powell 1994) as implemented in the nlopt library (Steven G. Johnson, The NLopt nonlinear-optimization package) to maximize the log-likelihood values calculated by the Forward algorithm. Convergence was attained in a maximum of 1000 evaluations and the log-likelihood maximization accuracy was set to 10^-4^. To test for convergence to local maxima, we ran the algorithm twice with different starting points and used the parameters of the run with the highest likelihood to run the re-estimation algorithm a third time starting with those parameters. All three runs gave similar results on all chromosomes.

### Post-processing

The HMM was executed independently on all chromosomes for both Denisova and Neandertal and using the African-American and DeCode recombination maps. An external region was defined as a stretch of high posterior probabilities (p ≥ 0.7) for the extended lineage sorting state that was uninterrupted by sites with a low probability (p ≤ 0.1). The two cutoffs on the posterior probabilities were determined by simulating sequences with positive selection (s=0.005, 500kya, see below). Sites that were simulated external in both Archaics were labeled as 1 and the remaining sites as 0. The HMM was then run on the simulations. By running a grid-search over possible cutoffs (step-sizes of 0.05 for the two parameters) and labeling the HMM output accordingly, we identified the set of chosen parameters by minimizing the root mean square error 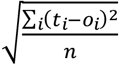 with n the number of labelled sites, t_i_ the true label and o_i_ the observed label.

### Simulations

We simulated sequences using a model of recent human demography to test the performance of our HMM under different scenarios of neutral evolution, positive selection or background selection. Each simulation consisted of one chimpanzee chromosome, one chromosome from each archaic hominin and 370 human chromosomes, matching the 185 Luhya and Yoruba individuals used in our analysis. For all simulations in this study, a constant mutation rate of 1.45×10^−8^ bp^−1^·generation^−1^, a constant recombination rate of 1cM.Mb^−1^.generation^−1^ and a generation time of 29 years were assumed. We used estimates of population sizes from (Yang et al. 2014) and population split estimates from (Prüfer et al. 2014) as parameters for the simulated demography (Supplemental Information 1 and 2). Neutral simulations with these parameters using the coalescent simulator scrm (Staab et al. 2014) give a good match to our observed data when plotting derived allele frequency in modern humans against the proportion of derived alleles in the outgroup (Supplemental Figure S8).

We generated a total of 100 loci of 1Mb-long sequences under neutrality to investigate the accuracy of labeling external and internal regions using our HMM. To evaluate the length of external regions expected under neutrality for the chromosome X, we simulated 100 loci of 1Mb-long sequences under the demographic model shown in Supplemental Information 1 with the exception that all effective population sizes were reduced to 75% of the original value. To evaluate the accuracy of parameter estimation, we additionally simulated splits of two populations (including an out-group individual) with a constant population size and different split times ranging from 400ky to 1My (step-size of 50ky). For each condition, we generated 25 sets of 10 Mb each. In an additional set of 100 loci of 1Mb, we introduced random errors by changing the state of the archaic allele with different rates in order to assess our error estimates.

To assess our power to detect events of positive selection, we explored selection coefficients ranging from 0.0005 to 0.1 and different times for the occurrence of the selected allele (every 100ky from 200kya to 600kya) using the coalescent simulator msms (Ewing and Hermisson 2010). The selected mutation was introduced in the middle of the sequence and we assumed an additive effect of the selected mutation (i.e. the homozygous genotype has twice the advantage stated by the selection coefficient). We performed 2000 simulations of 100kb-long loci for which all demographic parameters match our neutral simulations as described above. We used the –SForceKeep switch to drop the simulation if the selected mutation was lost. As 100kb loci are too short to make reliable parameter inferences, we concatenated our simulated sequences, intermittently combining them with 1Mb-long neutral loci from the previous simulations to limit the extent of the sequence affected by positive selection.

To explore the power over different settings of divergence, we simulated a simple demographic model with constant population size and varying degrees of divergence between two populations (see Supplemental Table S8 and S9 for further details).

We investigated how background selection affects lineage sorting in and around a conserved region by performing forward in time simulations using SLiM (Messer 2013). The simulated locus of 500kb length contained a conserved region resembling an ‘average’ human gene (see pg. 19 of the documentary accompanying SLiM (Messer 2013)) and covered 100kb (20%) of the simulated locus. Mutations in the conserved region were assumed to be neutral (25%) or deleterious (75%), with the selection coefficients of the deleterious mutations drawn from a gamma distribution with mean s = −0.05 and shape parameter α = 0.2. The deleterious mutations were assumed to be partially recessive with dominance coefficient h = 0.1 for a set of 100 simulations. To explore the effect of the strength of selection on the results, we produced 2 other sets of 40 simulations each by varying the mean of the gamma distribution (s = -0.001 and -0.1).

### Age Comparison with other Scans for Selection

To compare our sweep screen with previous scans, we downloaded candidate regions from the 1000G positive selection database (Pybus et al. 2014). Only candidates with a *P*-value lower than 0.001 were considered. We added to this set of regions the top reported regions from a HKA scan (Cagan et al. 2016). Allele age estimates were obtained from ARGweaver (Rasmussen et al. 2014).

Fst, iHS and XP-EHH are site-based statistics which localise sites that may have been selected (Sabeti et al. 2007; Malécot 1948; Voight et al. 2006; Wright 1951), whereas selective scans such as CLR, XP-CLR, Tajima’s D, Fay & Wu’s H and HKA identify candidate regions (Chen et al. 2010; Fay and Wu 2000; Hudson et al. 1987; Kim and Stephan 2002; Tajima 1989). In order to compare the age of the selection events, we assumed that the selected variant in candidate regions was the site with the highest frequency. We note that this procedure will underestimate the age of events if the true selected site reached fixation, as often expected for our method; the comparison is thus conservative.

### Annotations

We used the latest Ensembl gene annotation for hg19 (release 82) to identify protein-coding genes overlapping with our candidate regions. Based on this annotation, a regulatory region was defined as at least 5kb upstream and 1kb downstream of each gene. The regulatory region was extended until it reached a size of 1Mb or came within 5kb upstream or 1kb downstream of a neighboring gene. We additionally used a set of promoters and enhancers mapped by GenoSTAN in 127 cell types and tissues from the ENCODE and Roadmap Epigenomics projects (Zacher et al. 2016).

We used B-scores (McVicker et al. 2009) in hg19 coordinates constructed with UCSC’s liftover tool to evaluate the extent of background selection in our candidate regions. We also compared our candidate regions on the X chromosome with regions previously suggested to have experienced recurrent selective sweeps in apes (Dutheil et al. 2015; Nam et al. 2015). And, finally, we examined patterns of introgression in our candidate regions with two maps of Neandertal ancestry (Sankararaman et al. 2014; Vernot et al. 2014) and overlapped our regions with long deserts of Neandertal and Denisova ancestry from another recent study (Vernot et al. 2016).

To statistically test the overlap of our regions with these annotations, we permuted regions of similar physical sizes in the regions of the genome that passed our quality filters. Quality filtered regions that were smaller than the longest gap present in our candidate ELS regions were regarded as sufficiently short to not prohibit the placement of regions.

### Gene ontology and gene expression analysis

We defined genes that show tissue-specific expression levels using the Illumina BodyMap 2.0 RNA-seq data (Derrien et al. 2012), which contains expression data from 16 human tissues. We computed differential expression for all genes between a given tissue and all other tissues pooled using the DESeq package (Anders and Huber 2010) and genes were defined to be expressed in a tissue-specific manner when their expression levels were significantly higher (*P*-value < 0.05) in a given tissue compared to all other tissues. We tested for enrichment of candidate genes in the 16 sets of tissue-specifically expressed genes comparing to genes that were located outside of candidate regions using Fisher’s exact test. We calculated family-wise error rates for each tissue by randomly placing regions of sizes similar to the candidate regions in the genome. We repeated this process 1000 times, performed the same enrichment analysis as described above and counted how often any tissue in the randomized sets yields a smaller or equal *P*-value than the *P-*value observed in the candidate regions for a given tissue. This strategy corrects for the difference in length of genes expressed in specific tissues. We performed a similar analysis for the gene ontology analysis using func and the hypergeometric test (Prüfer et al. 2007), again comparing the genes associated with the candidate regions to a thousand sets of random regions to calculate family-wise error rates.

## DATA ACCESS

The software and input files used in this study have been made available through the website http://bioinf.eva.mpg.de/ELS/ and https://github.com/StephanePeyregne/ELS/.

## ACKNOWLEDGMENTS

We would like to thank Michael Lachmann for early discussions about the design of the study, Janet Kelso and Mark Stoneking for many useful comments on the manuscript, Svante Pääbo, Udo Stenzel, Fernando Racimo, Amy Ko and Adam Siepel for helpful discussions, Julien Peyrégne for help in implementing the software, and Matthias Ongyerth, Manjusha Chintalapati, Steffi Grote and Christoph Theunert for help with earlier analysis. This research was funded by the Max Planck Society and the Paul G. Allen Family foundation.

## AUTHOR CONTRIBUTIONS

SP implemented the method. SP, MJB and MD analyzed data. SP, MJB, MD and KP interpreted the results. KP designed the study. SP and KP wrote the manuscript with input from all authors.

